# *Fusobacterium nucleatum* metabolically integrates commensals and pathogens in oral biofilms

**DOI:** 10.1101/2021.06.23.449328

**Authors:** Akito Sakanaka, Masae Kuboniwa, Shuichi Shimma, Samar A. Alghamdi, Shota Mayumi, Richard J. Lamont, Eiichiro Fukusaki, Atsuo Amano

## Abstract

*Fusobacterium nucleatum* is a common constituent of the oral microbiota in both periodontal health and disease. Previously, we discovered ornithine cross-feeding between *F. nucleatum* and *Streptococcus gordonii*, where *S. gordonii* secretes ornithine via an arginine-ornithine antiporter (ArcD), which in turn supports the growth and biofilm development of *F. nucleatum*; however, broader metabolic aspects of *F. nucleatum* within polymicrobial communities and their impact on periodontal pathogenesis have not been addressed. Here, we show that when co-cultured with *S. gordonii*, *F. nucleatum* increased amino acid availability to enhance the production of butyrate and putrescine, a polyamine produced by ornithine decarboxylation. Co-culture with *Veillonella parvula*, another common inhabitant of the oral microbiota, also increased lysine availability, promoting cadaverine production by *F. nucleatum*. We confirmed that ArcD-dependent ornithine excretion by *S. gordonii* results in synergistic putrescine production, and mass spectrometry imaging revealed this metabolic capability creates a putrescine-rich microenvironment inside *F. nucleatum* biofilms. We further demonstrated that polyamines caused significant changes in the biofilm phenotype of a periodontal pathogen, *Porphyromonas gingivalis*, with putrescine being a potent stimulator of biofilm development and dispersal, and confirmed that *F. nucleatum*-mediated conversion of ornithine to putrescine enhances biofilm formation by *P. gingivalis*. Lastly, analysis of plaque samples revealed cooccurrence of *P. gingivalis* with genetic modules for putrescine production by *S. gordonii* and *F. nucleatum*. Overall, our results highlight the ability of *F. nucleatum* to induce synergistic polyamine production within multi-species consortia, and provide insight into how the trophic web in oral biofilm ecosystems can eventually shape disease-associated communities.

**Significance Statement:** Periodontitis is caused by the pathogenic transition of subgingival microbiota ecosystems, which is accompanied by alterations to microbiome functions including metabolic systems and the establishment of metabolic cross-feeding. While *Fusobacterium nucleatum* is a major constituent of the periodontal microbiota, its metabolic integration within polymicrobial communities and the impact on periodontal pathogenesis are poorly understood. Here, we report that amino acids supplied by other commensal bacteria induce polyamine production by *F. nucleatum*, creating polyamine-rich microenvironments. We further show that this trophic web results in enhancement of biofilm formation and dispersal of a periodontal pathogen, *Porphyromonas gingivalis*. This work provides mechanistic insight into how cooperative metabolism within oral biofilms can tip the balance toward periodontitis.

## Introduction

Periodontitis is a multifactorial chronic disease with diverse phenotypes often characterized by inflammatory destruction of periodontal tissues (1). The risk and severity of periodontitis are attributed to a dysbiotic transition in the community of microbes residing in the subgingival biofilm (2). In this process, *Porphyromonas gingivalis* plays a central role, although recent studies suggest that colonization of *P. gingivalis* does not necessarily elicit disease, and that full virulence requires the presence of the commensal microbiota, highlighting the importance of polymicrobial synergy in the disease etiology (3). Notably, a recent metatranscriptomic analysis of subgingival plaque from periodontitis patients showed highly conserved metabolic profiles, even though substantial microbiome variation was observed (4). This finding suggests that the transition between periodontal health and disease is more correlated with a shift in metabolic function of the community as a whole, rather than with the presence of individual taxa, drawing attention to metabolic aspects of microbial communities in periodontal pathogenesis.

Metabolic cross-feeding is one of the key factors directing the establishment of a community and the metabolism therein (5). A subset of oral streptococci engages in cross-feeding interactions with other community members that often result in elevated pathogenicity of microbial communities (6). A well-known example is lactate cross-feeding from *Streptococcus gordonii* to lactate-utilizing bacteria, such as *Veillonella parvula* and *Aggregatibacter actinomycetemcomitans*, where *S. gordonii* releases lactate as an end-product of glucose metabolism, thus allowing complementary utilization of available glucose and promoting fitness of these organisms in the community (7, 8). *S. gordonii* also impacts the pathogenicity of *P. gingivalis* through the secretion of para-aminobenzoic acid, which promotes in vivo fitness and colonization of *P. gingivalis*, albeit with diminished virulence (9). Given recent bioinformatic research showing that the oral microbiome can produce an enormous number of small metabolites that may influence oral pathophysiology (10), many more metabolic interactions between oral microbes likely remain to be discovered.

*F. nucleatum* is a common constituent of the oral microbiota, and has been implicated in both periodontal health and disease due to its frequent detection in subgingival plaque samples of both healthy and diseased sites (11–13). While this species is well known for its organizing role in oral biofilms through the expression of multiple adhesins, whereby it can direct the spatial relationships among early and later colonizers (14), metabolic aspects of interspecies interactions between *F. nucleatum* and other community members remain relatively unknown. Earlier studies showed that *F. nucleatum* supports the growth of *P. gingivalis* by rendering the microenvironment alkaline and less-oxidative (15). *F. nucleatum* has a preference for peptides and amino acids, and produces butyrate and ammonia as end-products of the fermentation pathways, starting mainly from glutamate and lysine (16, 17). The aforementioned metatranscriptomic analyses showed that despite nearly the same abundance of *F. nucleatum* between healthy and periodontitis samples, its metabolism is markedly changed under those two conditions (4). Considering that *F. nucleatum* is also strongly linked to serious systemic conditions such as adverse pregnancy outcomes and colorectal cancer (13), it is important to improve our basic understanding of the metabolic properties of *F. nucleatum* within polymicrobial communities.

Recently, we identified a novel metabolic interaction between *S. gordonii* and *F. nucleatum* (18), starting from the metabolism of arginine by *S. gordonii* as a substrate in the arginine deiminase system (ADS), through which arginine is converted to ornithine with concomitant production of ammonia and ATP. An arginine-ornithine antiporter of *S. gordonii*, ArcD, then excretes ornithine as a metabolic by-product of the ADS, which in turn enhances the growth and biofilm development of *F. nucleatum*. However, it is unknown how ornithine influences *F. nucleatum* metabolism and what consequences this interaction has for disease etiology. Therefore, in this study, we set out to further dissect the metabolic interactions mediated by *F. nucleatum* within multi-species consortia, and determine whether the engagement of *F. nucleatum* in metabolic interactions in oral biofilms can impact the potential pathogenicity of the microbial community. By using a synthetic community model to assess the metabolic changes of *F. nucleatum* and its microenvironment when co-cultured with *S. gordonii* and/or *V. parvula*, we show that the presence of partner species alters the amino acid metabolism in *F. nucleatum*, inducing production of butyrate and polyamines. We also demonstrate that putrescine production by *F. nucleatum* depends on ornithine cross-feeding via ArcD of *S. gordonii*. We further show that polyamines can modulate the biofilm phenotypes of *P. gingivalis*, with putrescine being a potent stimulator of biofilm formation and dispersal. This study has thus uncovered an emerging role of *F. nucleatum* as a metabolic bridge to relay the metabolic flow between initial and late colonizers, thereby creating favorable conditions for the outgrowth and spread of *P. gingivalis*.

## Results

### Distinct metabolic profiles in *F. nucleatum* co-cultured with *S. gordonii* and/or *V. parvula*

We performed untargeted analysis of the intra- and extracellular metabolite changes in *F. nucleatum* when co-cultured with *S. gordonii* and/or *V. parvula*. To facilitate metabolite exchange between different species and focus upon metabolic aspects of interspecies interactions, we used Transwell assays, which physically separate bacterial populations but allows for metabolite exchange via a shared medium reservoir. The system was anaerobically incubated in triplicate for 6 h in chemically defined medium (CDM) without organic nitrogen sources. Overall, we identified 111 extracellular and 85 intracellular metabolites, 52 of which were shared intra- and extracellularly (Dataset S1 and S2).

In co-culture with *S. gordonii*, orthogonal projection to latent structures-discriminant analysis (OPLS-DA) revealed that the intracellular metabolic profile of *F. nucleatum* clustered distinctly from that of *F. nucleatum* alone (Fig. 1A, inset), with putrescine, a product of ornithine decarboxylation, and *N*-acetylornithine, a product of ornithine acetylation, being associated with, and increased by, the presence of *S. gordonii* (Fig. 1A). Furthermore, we noted that the presence of *S. gordonii* elevated the intracellular concentrations of amino acids (alanine and glutamate) and a dipeptide (alanylalanine). Additionally, 16 extracellular metabolites were found in increased concentration using a fold change cutoff of 2 and a *p*-value of 0.05 (Fig. 1B). These metabolites were dominated by amino acids (ornithine, alanine, etc.) and products of amino acid fermentation and decarboxylation (butyrate, *N*-acetylputrescine, etc.). In particular, the relative concentrations of ornithine, alanine and butyrate were markedly increased in co-culture supernatants by 24.7-, 15.5-, and 9.4-fold, respectively. Further tests of *S. gordonii* mono-cultures confirmed that *S. gordonii* released all amino acids described here, some of which surpassed the levels during co-culture (e.g., ornithine, alanine, alanylalanine), suggesting net uptake of these metabolites by *F. nucleatum* (Fig. 1C). In contrast, fermented and decarboxylated products were undetected in the supernatant of *S. gordonii* alone, reflecting the metabolic potential of *F. nucleatum* to enhance production of these compounds in the presence of *S. gordonii*.

**Figure 1.**
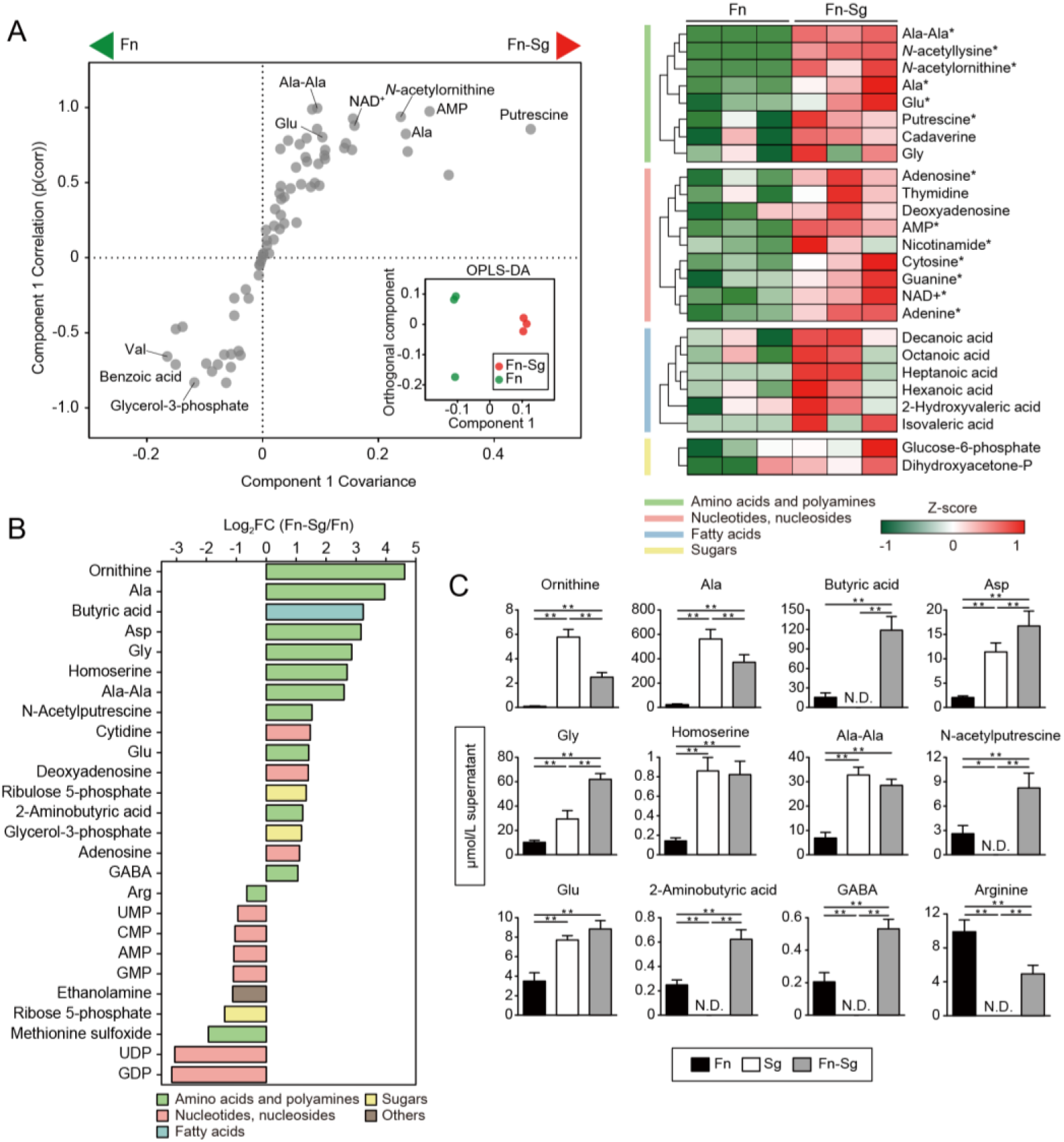
Intra- and extracellular metabolite changes of *F. nucleatum* co-cultured with *S. gordonii*. (A) Intracellular metabolite changes in *F. nucleatum* co-cultured with *S. gordonii*. 1.4×10^10^ cells of *F. nucleatum* were anaerobically cultured in CDM in the lower chamber of Transwell plates with membrane inserts, into which 1.4×10^10^ cells of *S. gordonii* in CDM or an equal volume of fresh CDM (as a control) were added. After 6 h, *F. nucleatum* cells were harvested and metabolic profiles were analyzed by CE-TOFMS. OPLS-DA S-plot and score plot (inset) are shown in the left panel, where metabolites towards the both sides of S-shape distribution show high model influence with high reliability; putrescine was among the most impacted metabolites in co-cultures. The right panel shows a clustered heatmap of intracellular metabolites with high reliability in the S-plot (p(corr) >0.6). Metabolite levels are displayed as Z scores, and asterisks denote significant differences in univariate methods. \**p* <0.05 (Mann-Whitney’s U test). (B) Extracellular metabolites displaying a concentration change in co-cultures as compared to mono-cultures (log_2_FC < −0.6, log_2_FC > 1 and *p* <0.05; Mann-Whitney’s U test). (C) Levels of the selected metabolites in spent media of co-cultures and each mono-culture, determined by UPLC. For this, the same procedures were repeated with the additional control of *S. gordonii* mono-cultures. Error bars correspond to standard deviations. \**p* <0.05 and \*\**p* <0.01 (one-way ANOVA with Tukey’s test).

In co-culture with *V. parvula*, OPLS-DA showed a discrete intracellular metabolite profile of *F. nucleatum*, in which lysine, dihydroxyacetone phosphate and thiamine were increased (Fig. 2A). Additionally, we found increased levels of seven extracellular metabolites in co-culture, four of which were products of amino acid fermentation and decarboxylation. In particular, cadaverine, a product of lysine decarboxylation, exhibited the most prominent change (Fig. 2B). Since cadaverine was undetected in the supernatants of *V. parvula* mono-cultures (Fig. 2C), *F. nucleatum* is likely to produce cadaverine by utilizing lysine released by *V. parvula*. In co-culture with a mixed population of *S. gordonii* and *V. parvula*, we observed an additive effect of these two species on the intra- and extracellular metabolic profiles of *F. nucleatum* (Fig. S1).

**Figure 2.**
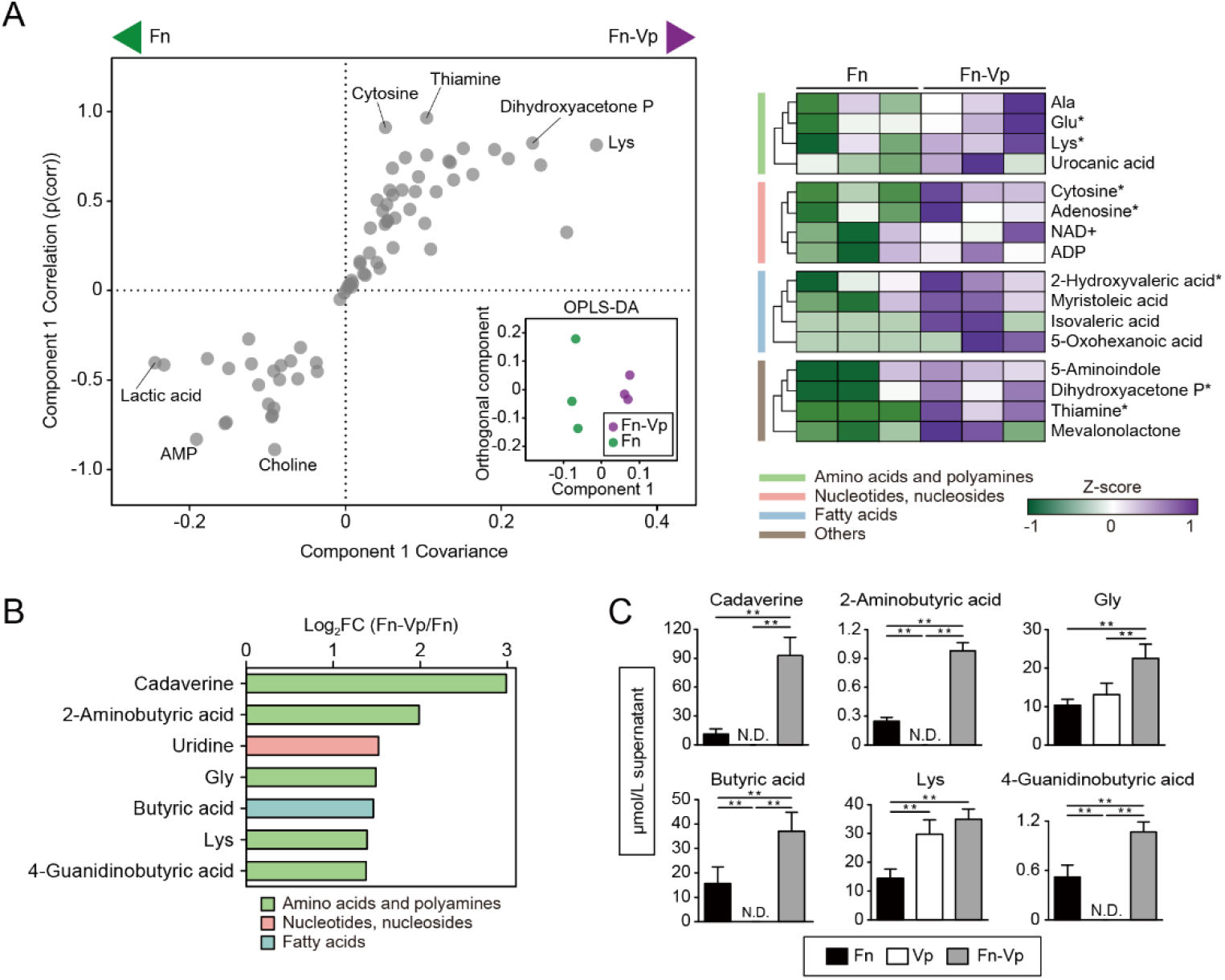
Intra- and extracellular metabolite changes of *F. nucleatum* co-cultured with *V. parvula*. (A) Intracellular metabolite changes in *F. nucleatum* co-cultured with *V. parvula*. OPLS-DA score plot (inset) and S-plot (left panel) show that lysine and thiamine were among the most impacted metabolites in co-cultures. The right panel shows a clustered heatmap of intracellular metabolites with high reliability in the S-plot (p(corr) >0.6). \**p* <0.05 (Mann-Whitney’s U test). (B) Extracellular metabolites displaying a concentration change in co-cultures as compared to mono-cultures (log_2_FC < −0.6, log_2_FC > 1 and *p* <0.05; Mann-Whitney’s U test). (C) Levels of the selected metabolites in spent media of co-cultures and each mono-culture, determined by UPLC. Error bars correspond to standard deviations. \**p* <0.05 and \*\**p* <0.01 (one-way ANOVA with Tukey’s test).

### Upregulation of butyrate fermentation and polyamine production by *F. nucleatum* in co-culture

To gain further insight into these metabolic interactions, we assessed transcriptional changes in related genes of *F. nucleatum* using real-time RT-PCR under the same culture conditions as those of the metabolomics assays. In co-culture with *S. gordonii*, we observed an upregulation of a cluster of genes encoding critical enzymes for butyrate production, including FN0202-0204, which is located in the 2-hydroxyglutarate pathway and links butyrate to glutamate (Fig. 3A). The same trend was observed when *F. nucleatum* was co-cultured with mixtures of *S. gordonii* and *V. parvula*, while the presence of *V. parvula* had a minor effect on the transcriptional activation of butyrate fermentation pathways, with only two enzymes of the butyrate production from lysine pathway upregulated. Since amino acid fermentation contributes to energy generation in anaerobic bacteria, we measured the ATP levels in *F. nucleatum* cells in co-culture. We found a 1.87-fold increase in ATP levels per cell in *F. nucleatum* co-cultured with *S. gordonii* (Fig. 3B). Collectively, these data indicate that coexistence with *S. gordonii* facilitates butyrate production, especially from glutamate, by *F. nucleatum*, thereby promoting ATP generation.

**Figure 3.**
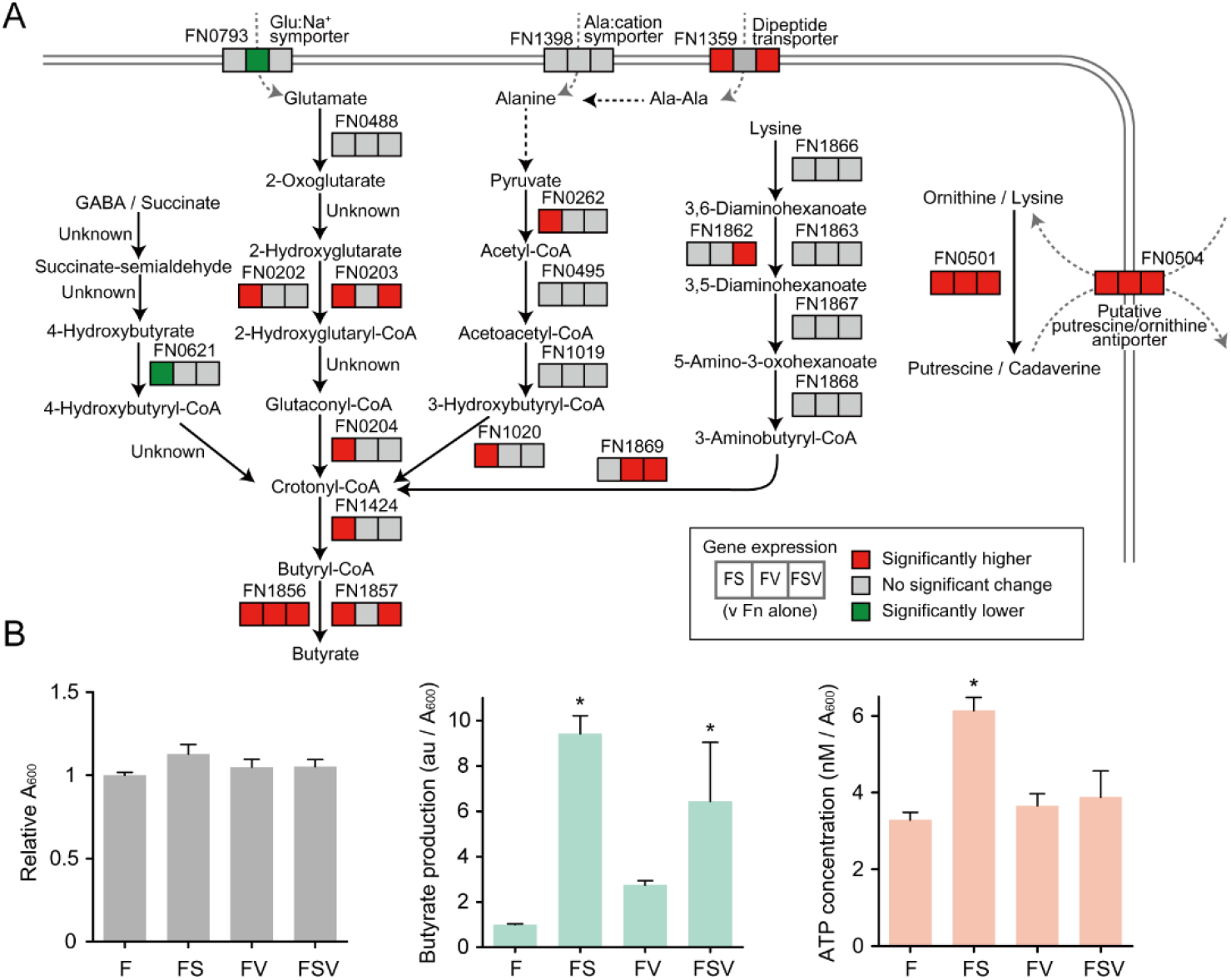
Upregulation of butyrate fermentation and polyamine production by *F. nucleatum* in co-culture. (A) Transcriptional changes of selected genes involved in the production of butyrate and polyamines by *F. nucleatum* when co-cultured with *S. gordonii* or *V. parvula* individually or in combination. Transcripts were extracted from *F. nucleatum* cells following the same culture conditions as those used for metabolomic assays. 16S rRNA was used for normalization. Statistical differences were analyzed using a one-way ANOVA with post hoc paired comparisons conducted with Dunnett’s test (*p* <0.05). Red denotes significantly increased levels (>1.5-fold change), green decreased levels (<0.65-fold change) and gray no significant changes. (B) Relative production of butyrate and ATP by *F. nucleatum* in each condition. The left panel shows relative absorbance changes in *F. nucleatum* biomass after 6 h of incubation in each condition. In this assay, biofilm cells were also retrieved to comprise a total biomass. Bars are representative of three independent experiments and presented as the mean with SD of three biological replicates. The center panel shows the A_600_-adjusted abundance (mean ± SD) of butyrate in culture supernatants from the metabolomics dataset. The right panel shows the A_600_-adjusted ATP concentration in *F. nucleatum* cells after 6 h of incubation. F: *F. nucleatum* alone; FS: *F. nucleatum* and *S. gordonii*; FV: *F. nucleatum* and *V. parvula*; FSV: *F. nucleatum* with *S. gordonii* and *V. parvula*. Bars show the mean with SD of a representative experiment of five biological replicates. *, *p* < 0.05 (versus *F. nucleatum* alone, calculated using ANOVA with Dunnett’s test).

Putrescine and cadaverine are most commonly produced by the decarboxylation of ornithine and lysine, reactions catalyzed by ornithine decarboxylase (encoded by *speC*; Enzyme Commission number, E.C. 4.1.1.17) and lysine decarboxylase (*cadA*; E.C. 4.1.1.18), respectively. A newly reannotated database of *Fusobacterium* genomes shows the presence of a gene containing the domain of ornithine and lysine decarboxylases in *F. nucleatum* ATCC25586 (FN0501), which shows high similarity to the sequences of both the *speC* and *cadA* genes of *Escherichia coli* (19, 20). We found that the relative transcriptional level of FN0501 was elevated greater than 20-fold in all pairs of co-cultures in our assays (Fig. 3A). Furthermore, *F. nucleatum* possesses a gene (FN0504) which shows high similarity to the putrescine/ornithine antiporter of *E. coli* (21), and this gene was also transcribed at a significantly increased level in the presence of *S. gordonii* and/or *V. parvula*.

Collectively, these results suggest that the presence of *S. gordonii* and *V. parvula* increases amino acid availability for *F. nucleatum*, resulting in enhanced production of fermented and decarboxylated metabolites. Notably, *F. nucleatum* is likely to produce putrescine and cadaverine via decarboxylation of ornithine and lysine released by *S. gordonii* and *V. parvula*, respectively.

### Commensal-triggered polyamine production by *F. nucleatum*

We next tested whether putrescine production results from ArcD-dependent excretion of ornithine by *S. gordonii*. We incubated mixtures of *F. nucleatum* with WT or *S. gordonii* Δ*arcD* as well as mono-cultures of each strain in CDM containing 10 mM arginine, and quantified concentrations of arginine, ornithine and putrescine in the culture supernatants. After 24 h, WT *S. gordonii* consumed arginine completely and released 8.26 mM ornithine, but was unable to produce putrescine by itself (Fig. 4A). Similarly, *F. nucleatum* alone failed to utilize arginine or to produce putrescine and ornithine. In contrast, co-cultures of *F. nucleatum* and WT *S. gordonii* depleted arginine and released 3.55 mM ornithine and 2.94 mM putrescine, which together with ammonia produced via ADS, allowed for maintenance of neutral pH in culture supernatants (Fig. 4A, far right). Lack of ArcD suppressed not only arginine uptake and ornithine release by *S. gordonii*, as demonstrated in our previous work (18), but also putrescine production in co-cultures. Next, we used spent medium from mono-cultures of each organism to culture the other, and quantified arginine, ornithine and putrescine in the culture supernatants. *S. gordonii* depleted 10 mM arginine and released 7.27 mM ornithine during 12 h-cultivation (Fig. 4B). When *F. nucleatum* was cultured using these supernatants, ornithine decreased from 7.27 to 4.93 mM, while putrescine increased from 0 to 2.43 mM. In contrast, arginine remained intact when *F. nucleatum* was initially cultured, and cultivation of *S. gordonii* using these supernatants failed to produce putrescine. Collectively, these results indicate that production of putrescine by *F. nucleatum* depends on release of ornithine from *S. gordonii* as a metabolic by-product of the ADS.

**Figure 4.**
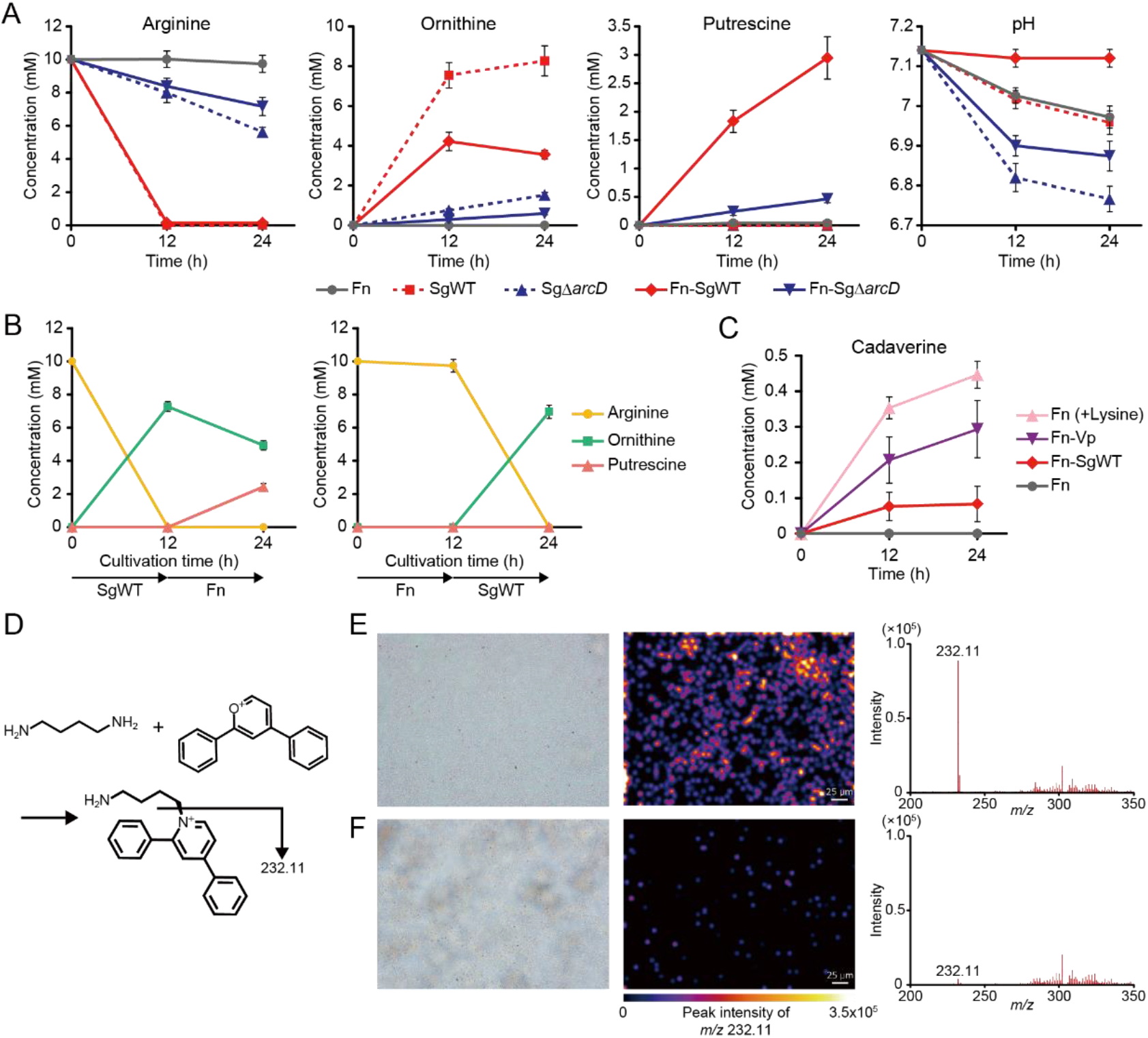
Commensal-triggered polyamine production by *F. nucleatum*. (A) Extracellular concentrations of arginine, ornithine and putrescine in CDM containing 10 mM arginine incubated anaerobically for 12 and 24 h were determined by UPLC after bacterial cells were removed. Extracellular pH changes were also shown. (B) Shifts in the extracellular concentrations of arginine, ornithine and putrescine in CDM containing 10 mM arginine incubated initially with *S. gordonii* or *F. nucleatum* for 12 h then with its counterpart for additional 12 h. (C) Changes in cadaverine concentrations were determined in supernatants of the designated cultures. Data are shown as the means with SDs of a representative experiment of three biological replicates. (D) Schematic of putrescine imaging. Using 2,4-diphenyl-pyranylium tetrafluoroborate (DPP-TFB), on-tissue derivatization was performed, and the distribution of putrescine (target *m/z* 232.11) was visualized through matrix-assisted laser desorption/ionization mass spectrometry imaging (MALDI-MSI). Shown are optical images and imaging results of biofilms formed on indium-tin-oxide (ITO)-coated glass slides by immersion for 24 h in *F. nucleatum* mono-cultures developed in PBS (E) with or (F) without 10 mM ornithine. Color brightness corresponds to concentration of putrescine.

We then incubated axenic cultures of *F. nucleatum* in the presence of 10 mM lysine or mixed cultures with *V. parvula* or WT *S. gordonii* in CDM and quantified cadaverine in the culture supernatants. After 24 h, 0.44 mM cadaverine was produced in the lysine-incubated axenic cultures, while co-cultures with *V. parvula* produced 0.29 mM cadaverine, suggesting synergistic cadaverine production via lysine cross-feeding between these species (Fig. 4C).

To further validate the ability of *F. nucleatum* to metabolize ornithine to putrescine and to evaluate its consequence on biofilm microenvironments, we employed matrix-assisted laser desorption/ionization mass spectrometry imaging (MALDI-MSI) and quantitatively visualized the spatial distribution of putrescine within *F. nucleatum* biofilms formed on glass slides treated with or without 10 mM ornithine. Illustrations of ion signals for putrescine revealed an abundance of putrescine deposited within the biofilm treated with ornithine (Fig. 4E and F), providing evidence that *F. nucleatum* can alter the metabolic landscape in the biofilm by creating a putrescine-rich microenvironment.

### Polyamines can enhance the pathogenic potential of *P. gingivalis* via modulation of the biofilm phenotype

The results presented above indicate that metabolic interactions among oral commensals can induce polyamine production by *F. nucleatum*. To explore the consequences of these interactions on the development of disease-associated communities, we assessed the effects of polyamines on the biofilm phenotype of a periodontal pathogen, *P. gingivalis*. For this experiment, we used the most widely distributed bioactive polyamines (putrescine, spermidine, spermine and cadaverine), whose release from *F. nucleatum* was also confirmed in the Transwell assays (Dataset S1). We incubated preformed-*P. gingivalis* biofilms anaerobically with each polyamine for 12 h. After staining with Live/Dead reagent and 3 h-additional incubation with each polyamine, the amount and viability of the biofilm and of planktonic cells were evaluated using confocal laser scanning microscope (CLSM). Analysis of the biofilm structures showed the stimulatory effects of putrescine, cadaverine (*p* <0.01) and spermidine (*p* <0.05) on biofilm development; in particular, exogenous putrescine caused the greatest increase in not only the viable attached biofilms but also the viable planktonic biomass, which had dispersed from the post-stained biofilms (Fig. 5A and C). In contrast, cadaverine exhibited an opposite trend in this regard, producing more rigid biofilms with less suspended planktonic cells (Fig. 5B). These results suggested that these polyamines have discrete effects on the biofilm phenotype of *P. gingivalis*. Furthermore, we observed a dose-dependent effect of putrescine on the amounts of biofilms as well as dispersed cells of *P. gingivalis* (Fig. 5E), suggesting a potent stimulatory effect on both biofilm formation and dispersal. To test whether the observed effects of polyamines were specific to *P. gingivalis*, we performed additional controls using other bacteria. Unlike *P. gingivalis*, *S. gordonii* was relatively insensitive to polyamines, with its dispersal behavior repressed, and its biofilm formation was promoted only by spermidine (Fig. 5D). In contrast, spermine exhibited biofilm disruptive activity against *F. nucleatum*, and cadaverine increased both the biofilm and planktonic biomass of *V. parvula* (Fig. S2). These results indicate that polyamines have a diversity of physiological functions in different oral bacteria. Finally, we observed the response of *P. gingivalis* to pH-adjusted cell-free supernatants from co-cultures of *F. nucleatum* and WT/ *S. gordonii* Δ*arcD*. We found that cell-free supernatants from co-cultures of *F. nucleatum* and WT *S. gordonii* significantly enhanced biofilm formation by *P. gingivalis* (Fig. 5F). Additionally, mixed biofilm experiments showed that metabolism of ornithine by *F. nucleatum* produces a synergistic effect on *P. gingivalis* biofilm growth (Fig. 5G and H) Together, these data showed that polyamines produced by *F. nucleatum* can impact the biofilm phenotypes of *P. gingivalis*, with putrescine being a potent stimulator of biofilm development and dispersal.

**Figure 5.**
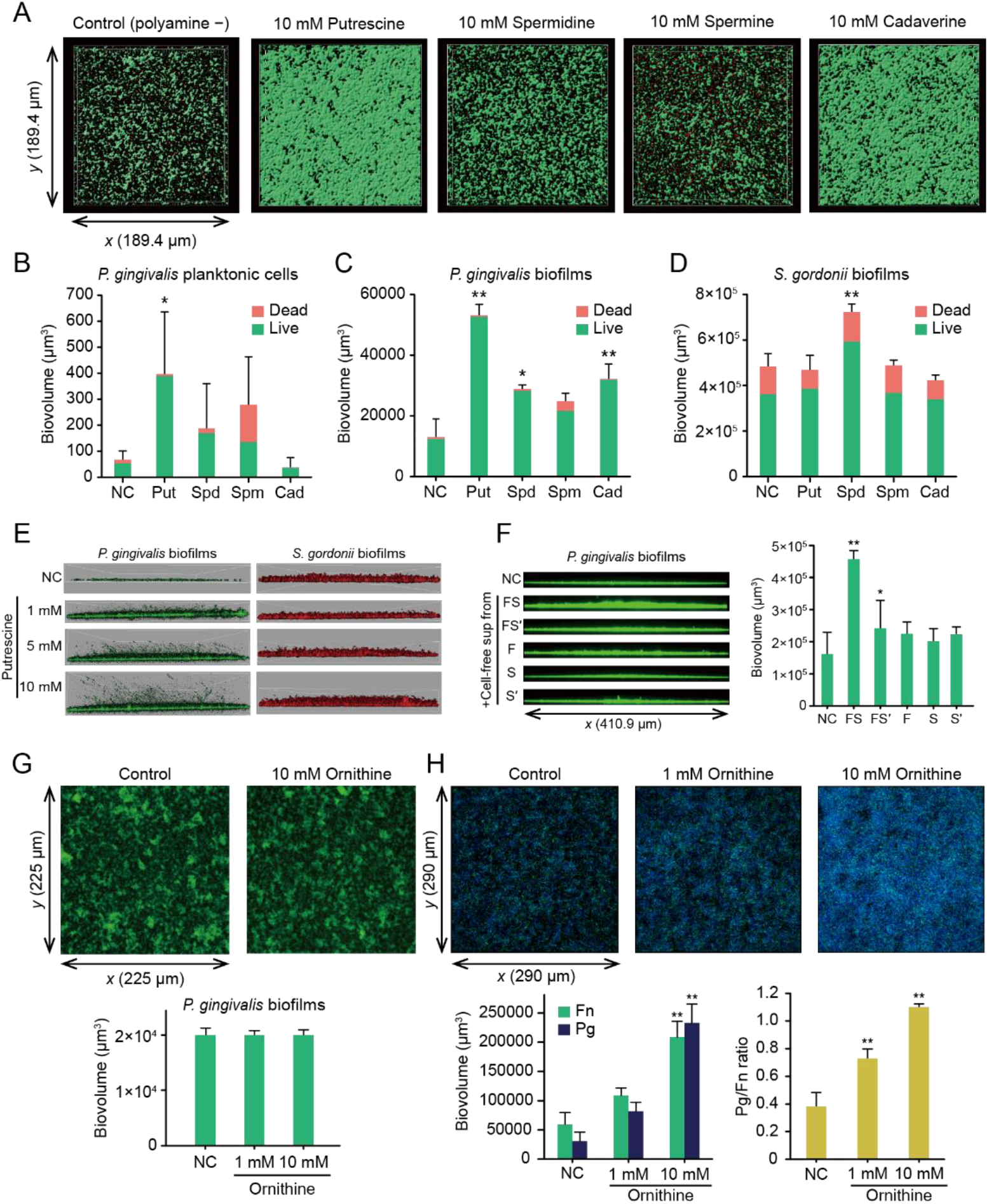
Effects of exogenous polyamines on biofilm growth and dispersal of *P. gingivalis*. (A) Preformed *P. gingivalis* biofilms were treated anaerobically with PBS containing each polyamine for 12 h and then stained with Live/Dead dyes. A series of optical fluorescence x-y sections were collected by confocal microscopy. Images are representative of three independent experiments. Biovolumes of dispersed planktonic cells (B) and biofilm cells (C) were measured with the Imaris Isosurface function after reconstructing three-dimensional images by applying an isosurface over Live/Dead-stained biomass separately per color (green/red). (D) The effects of each polyamine on *S. gordonii* biofilms were examined as a control, following the same method. Data are representative of three independent experiments and presented as the mean with SD of eight random fields from one experiment. \**p* <0.05, \*\**p* <0.01 compared with the control using ANOVA with Dunnett’s test. (E) Representative images of putrescine-treated biofilm microstructures of *P. gingivalis* and *S. gordonii*, which were stained with FITC and HI, respectively, at the start of the experiment. (F) *P. gingivalis* biofilms were formed in PBS containing 50% cell-free pH-adjusted supernatants of each culture incubated anaerobically for 24 h. FS denotes cell-free supernatants of mixed cultures of *F. nucleatum* and WT *S. gordonii*. FSʹ denotes those of *F. nucleatum* and *S. gordonii* Δ*arcD*, while F, S and Sʹ denotes those of mono-cultures of *F. nucleatum*, WT and *S. gordonii* Δ*arcD*, respectively. \**p* <0.05, compared with the control using ANOVA with Dunnett’s test. (G) Effect of ornithine on *P. gingivalis* biofilms. FITC-stained *P. gingivalis* biofilms were formed in the presence of ornithine after 24 h of incubation. Representative images of biofilm architecture and biovolume of *P. gingivalis* are shown. (H) Effect of ornithine on *P. gingivalis* accumulation in *F. nucleatum* biofilms. FITC-stained *F. nucleatum* biofilms (green) were formed after 24 h of incubation, then gently washed with PBS and co-cultured for 24 h with DAPI-labelled *P. gingivalis* (blue) in the presence of ornithine. Representative images, biovolumes of each species and their ratios are shown. \*\**p* <0.01 compared with the control using ANOVA with Dunnett’s test.

### Cooccurrence of *P. gingivalis* with genetic modules for putrescine production by *S. gordonii* and *F. nucleatum* in plaque samples

To test the applicability of the results to the human oral cavity, we analyzed plaque samples from 102 systemically healthy individuals and investigated the relationship between the presence of *P. gingivalis* and the levels of the *arcD* gene of *S. gordonii*, and FN0501 of *F. nucleatum* using real-time PCR. We found that *P. gingivalis* was detected more frequently as periodontal health deteriorates (Fig. 6A). Furthermore, the *arcD* gene of *S. gordonii* exhibited a higher abundance in *P. gingivalis* positive samples (Fig. 6B), and a combination of *arcD* and FN0501 genes by logistic regression achieved areas under the curve of 0.76 for *P. gingivalis* detection, surpassing the discriminative performance of periodontal inflamed surface area, a numerical representation of periodontitis severity (22) (Fig. 6C). These data provide clinical evidence suggesting cooccurrence of *P. gingivalis* with genetic modules for putrescine production by *S. gordonii* and *F. nucleatum*. Based on these results, we propose a model of metabolic interactions within oral biofilms whereby ADS in *S. gordonii* facilitates putrescine production by *F. nucleatum* which could further promote the biofilm overgrowth and dispersal of *P. gingivalis* (Fig. 7A and B).

**Fig. 6.**
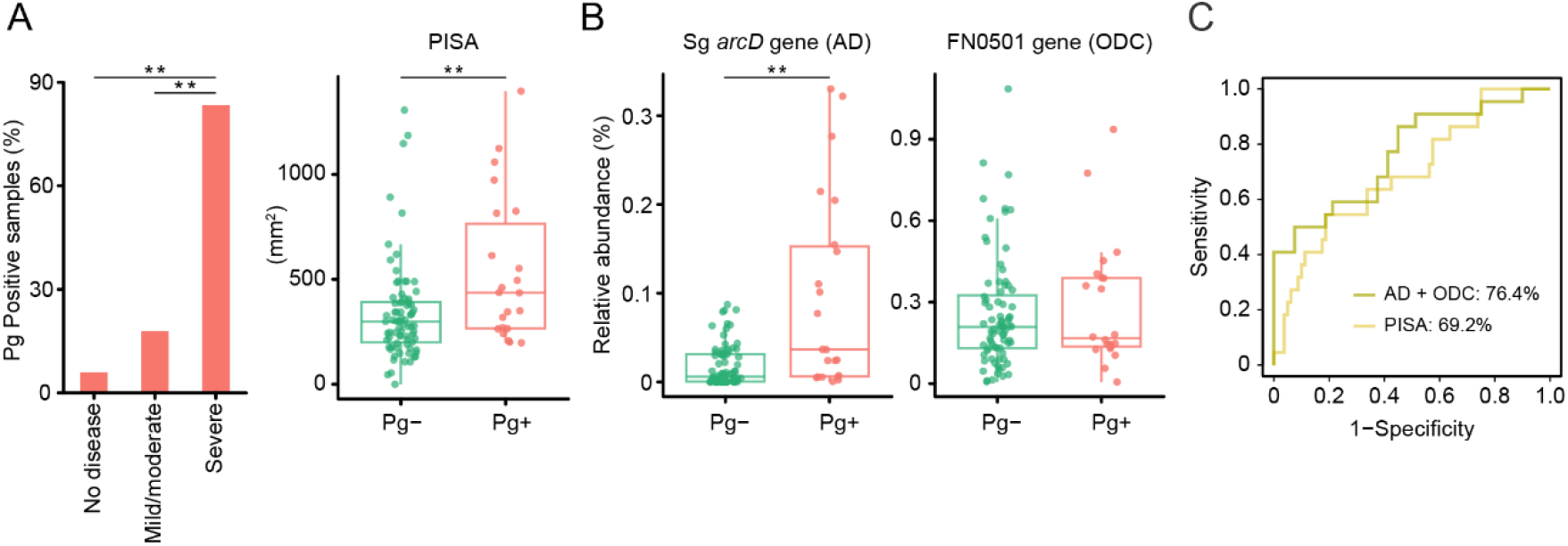
Cooccurrence of *P. gingivalis* with genetic modules for putrescine production by *S. gordonii* and *F. nucleatum* in 102 plaque samples. (A) Detection of *P. gingivalis* in supragingival biofilms in states of periodontal health, mild/moderate periodontitis and severe periodontitis (left). Difference in periodontal inflamed surface area (PISA), a numerical representation of periodontitis severity, between *P. gingivalis* positive and negative samples (right). \*\**p* <0.01 compared with “no disease” using chi-square test (left). \*\**p* <0.01 Mann-Whitney’s U test (right). (B) Difference in abundances of *S. gordonii arcD* gene and *F. nucleatum* FN0501 gene between *P. gingivalis* positive and negative samples. \*\**p* <0.01 Mann-Whitney’s U test. (C) ROC curves comparing the discriminative performance for *P. gingivalis* detection using logistic regression with *arcD* and FN0501 genes (olive), and PISA (yellow).

**Fig. 7.**
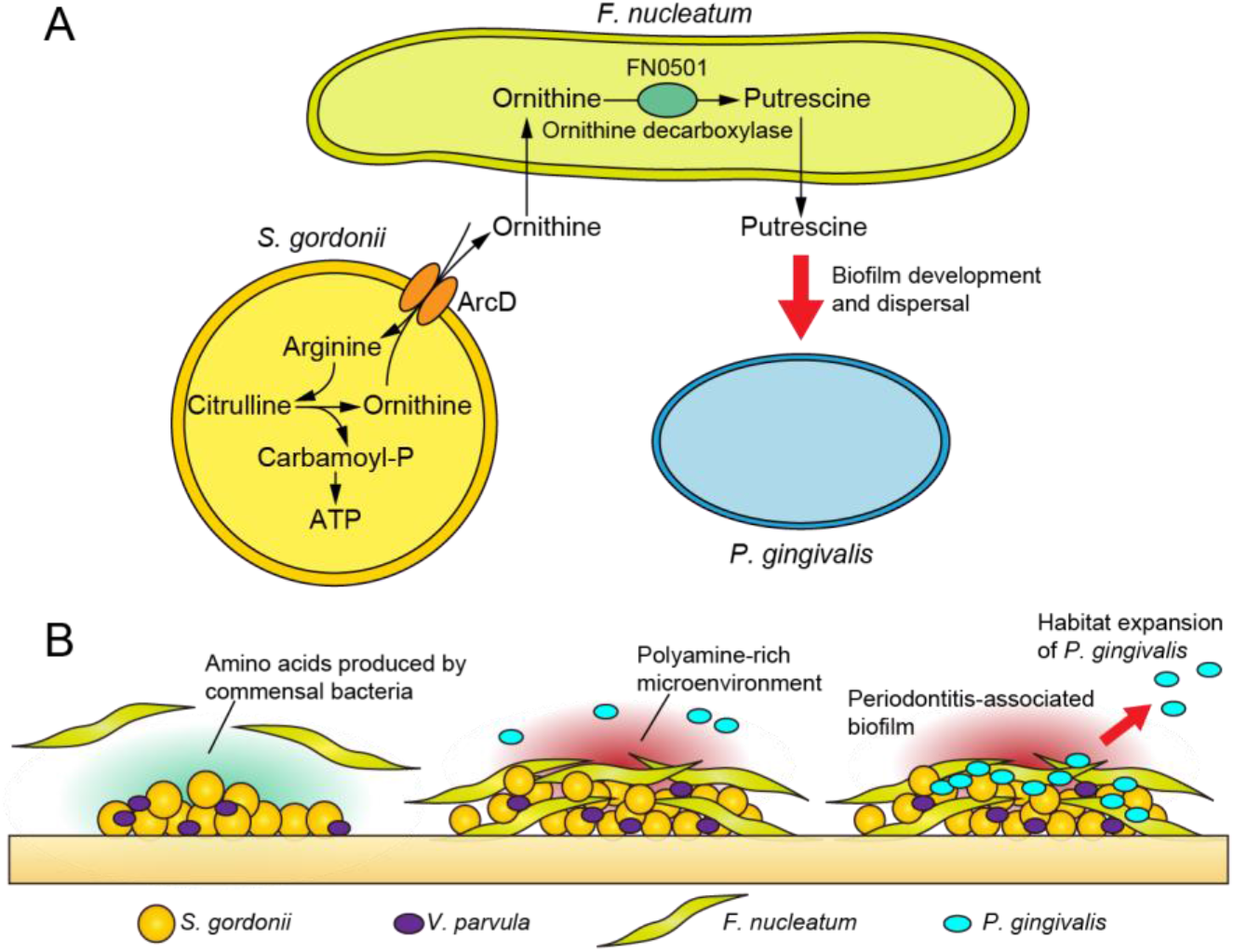
Proposed schematic model of polymicrobial metabolic synergy in the disease etiology. (A) Pathogenic cross-feedings among three key species. Arginine deiminase system in *S. gordonii* facilitates putrescine production by *F. nucleatum*, which could further promote the biofilm overgrowth and dispersal of *P. gingivalis*. (B) Model depicting metabolic integration by *F. nucleatum* within polymicrobial communities. Commensal-triggered polyamine production by *F. nucleatum* contributes to shaping the periodontitis-associated community.

## Discussion

Oral bacterial communities often exhibit synergistic pathogenesis via physical and metabolic interactions among community members, which perturbs host homeostasis and ultimately causes periodontitis (2). *F. nucleatum*, which is a major constituent of the periodontal microbiota, contributes to this process by connecting a diverse range of microbes and lending physical support to the biofilm structure (13). Here, we report that engagement of *F. nucleatum* in metabolic interactions with other commensal bacteria creates polyamine-rich microenvironments, which facilitate biofilm development and dispersal of *P. gingivalis*, thereby potentially impacting the pathogenicity of periodontitis.

Combining the results of intra- and extracellular metabolite changes in *F. nucleatum* from Transwell assays suggested possible engagement of some amino acids in cross-feeding interactions. Specifically, in addition to ornithine, consistent with our previous report (18), *F. nucleatum* acquired alanine, alanylalanine and glutamate released from *S. gordonii* and lysine from *V. parvula*, although it is unclear if these metabolites are released via leakage or specific transport. Amino acids are the main source of energy for *F. nucleatum*, but it does not possess a high level of endopeptidase activity (23), and previous studies propose that it takes advantage of amino acids and peptides available through interspecies interactions with proteolytic bacteria such as *P. gingivalis* (24, 25). Our results suggest that amino acids can also be supplied by oral commensals lacking proteolytic activity through a cross-feeding behavior. Indeed, emerging evidence suggests that amino acid cross-feeding is one of the main drivers of interspecies interactions in microbial communities (5, 26, 27). Recent experimental observations show that diverse microbial species secrete amino acids without fitness cost, which generates ample cross-feeding opportunities, and can be facilitated by anoxic conditions (28). In light of this, our results support the plausibility of widespread amino acid cross-feeding within both the supragingival and subgingival communities, which could underlie metabolic shifts during the transition from periodontal health to disease.

One of the most striking findings in this study was that *S. gordonii* and *F. nucleatum* interact cooperatively to produce putrescine from arginine through ornithine, and this trophic web results in alterations in *P. gingivalis* biofilm phenotypes. Conversion of ornithine to putrescine via decarboxylation consumes cytoplasmic protons and creates a proton motive force (29), offering an energetic advantage to *F. nucleatum*. Consumption of ornithine also helps maintain the ADS function and achieve a sustainable energy supply for *S. gordonii*. From ecological and evolutionary perspectives, therefore, this collaborative metabolism accomplished by the ADS of *S. gordonii* and ornithine decarboxylase of *F. nucleatum* would be favored by natural selection, since it allows for the efficient use of limited resources and confers fitness benefits to both species. Moreover, this sequential reaction eventually created an alkaline microenvironment via ADS-dependent ammonia and *F. nucleatum*-derived putrescine (Fig. 4A). Given the biofilm-stimulating effect of putrescine, these chemical and metabolic changes favor the survival of *P. gingivalis*, suggesting that this mutualistic interaction between these three species underlies the enhancement of community pathogenicity.

In addition, *F. nucleatum* has been implicated in colorectal cancer (13), and a recent imaging-based analysis of colon tumors has revealed an abundance of acetylated polyamines in colorectal biofilm samples (30). Although further studies are needed, our findings suggest that the contribution of this species to colon tumorigenesis could be attributed, at least partially, to the polyamine production system.

It should be noted that significant extracellular accumulation of putrescine was observed in polyamine production assays (Fig. 4) while not in Transwell assays (Fig. 1B). Given that polyamines have diverse cellular functions and their intracellular levels are strictly regulated in bacteria (31), one possible explanation is that putrescine secretion by *F. nucleatum* is likely due to metabolic overflow, where a certain amount of ornithine in the medium triggers overproduction of putrescine, inducing its secretion (32). We consider that the extracellular level of ornithine had yet to reach this amount in Transwell assays, where *F. nucleatum* and *S. gordonii* were incubated without arginine for only 6 h. Additional studies are required to elucidate the underlying mechanisms regulating production and secretion of putrescine in this organism, and we are addressing this question by targeting several polyamine transporters of *F. nucleatum*, including a putative polyamine ABC transporter and FN504. In addition, polyamine homeostasis is known to be maintained by acetylation of its substrates and products (33), and we detected some acetylated metabolites in *F. nucleatum* (e.g. *N*-acetylornithine, *N*-acetylputrescine, Fig. 1A), whose roles in polyamine metabolism have also to be investigated.

Putrescine is regarded as one of the most common polyamines in bacteria, and together with spermidine, its biosynthesis was found to be essential for the growth of many opportunistic pathogens, including *Pseudomonas aeruginosa* and *Campylobacter jejuni* (34, 35). Putrescine and spermidine are also required for biofilm formation by *Bacillus subtilis* and *Yersinia pestis* (36–38), although spermidine inhibits biofilm formation by some bacteria (39, 40). Here, we demonstrated that exogenous putrescine and cadaverine stimulate *P. gingivalis* biofilm development while producing different biofilm phenotypes; cadaverine yielded more rigid biofilms with less suspended cells, whereas putrescine thickened biofilms with more suspended cells. The distinct biofilm phenotypes may represent differences in biofilm developmental stages and suggest the potential of putrescine to accelerate the lifecycle of *P. gingivalis* biofilms, enabling both biofilm formation and dispersal. In fact, previous work showed that putrescine acts as an extracellular signal for swarming and is necessary for effective migration across agar surfaces in *Proteus mirabilis* (41). A recent multi-omics study showed that *P. gingivalis* strain 381 can surface translocate when sandwiched between two surfaces, and this dispersion-like behavior involves intracellular metabolic changes in the arginine and polyamine pathways, with citrulline and ornithine accumulation along with exhaustion of arginine and putrescine (42). Although the mechanistic details of the role of polyamines in *P. gingivalis* physiology are largely unknown and further studies will be necessary to gain a better understanding, putrescine seems to be a key signal for transforming physiology and accelerating the biofilm lifecycle of *P. gingivalis* to promote habitat expansion.

A number of studies using clinical samples have found the possible involvement of polyamines and related metabolites in the pathogenesis of periodontitis. A comparative metagenomics study using whole-genome shotgun sequencing revealed that a disease-associated microbiota exhibits metabolic functions that are largely absent in health, and those functions include polyamine uptake systems regulated by a putrescine transport ATP-binding protein (43). A metabolomic analysis of gingival crevicular fluids revealed significantly elevated levels of putrescine and cadaverine, as well as various amino acids including ornithine, in the subgingival crevice of periodontitis sites (44). Our previous metabolomic studies using saliva samples also showed that a disease-associated microbiota likely produces polyamines, including putrescine and cadaverine, which is reflected in the distinct salivary metabolomic landscapes of periodontitis patients (45, 46). Although the extent to which *F. nucleatum* dictates the enrichment of polyamine metabolism in periodontitis has yet to be determined and other community members may contribute to polyamine production, the data presented in this work add to the evidence that the transition from periodontal health to disease is linked to metabolic specialization, including polyamine metabolism, in subgingival microbial communities.

Although this study focused on a few oral bacteria to simulate metabolic cross-feeding during dental biofilm maturation, we acknowledge that oral biofilm ecosystems have food webs comprising many layers of complexity that fall outside the scope of our framework (47). In addition, the nature of metabolic interactions may be affected by the physical proximity of species and their structural organization, which are important features of biofilms (48). These limitations notwithstanding, this study provides new insights into how the trophic web in oral biofilm ecosystems impacts the process of dental biofilm maturation; specifically, ornithine cross-feeding by *S. gordonii* induces putrescine production by *F. nucleatum*, which can culminate in the overgrowth and habitat expansion of *P. gingivalis*. Our results reveal a new example of cooperative metabolism between oral bacteria that is unattainable without the sharing of metabolic pathways in multiple taxa, and shed light on the metabolic aspects of *F. nucleatum* in the context of the pathogenicity of microbial communities through metabolic communications within oral biofilms. Given the significant impact of polyamines on *P. gingivalis* phenotypes, future work will address the mechanisms by which polyamines affect the physiology of *P. gingivalis* and explore the possibility that assessing polyamine profiles in subgingival biofilms may yield a novel method for monitoring disease activity and eventually lead to disease prevention.

## Materials and Methods

### Bacterial strains and growth conditions

*F. nucleatum* subsp. *nucleatum* ATCC 25586, *P. gingivalis* ATCC 33277 and *V. parvula* JCM 12972 were grown statically at 37°C in an anaerobic chamber (Concept Plus; Ruskinn Technology, Bridgend, UK) containing 10% H_2_, 10% CO_2_, and 80% N_2_. Solid and liquid media used for growing each species are described in Supplementary Information. *S. gordonii* DL1 and its isogenic Δ*arcD* mutant (18) were grown statically in Todd-Hewitt broth (Becton, Dickinson and Company, Franklin Lakes, NJ, USA) under aerobic conditions (in 5% CO_2_ at 37°C), and erythromycin (5 mg/L) was used for selection. At the early-stationary phase, bacteria were harvested by centrifugation, washed twice with pre-reduced phosphate-buffered saline (PBS), and then used in the assays. For Transwell assays, bacteria were anaerobically cultured at 37°C in CDM containing inorganic salts, vitamins and 0.1% glucose (see Supplementary Materials and Methods for detailed composition).

### Transwell assay and metabolomic and transcriptional analyses

Synthetic communities were created by inoculating 1.4×10^10^ cells of *F. nucleatum* in CDM in the lower chamber of a 6-well Transwell system with 0.4-μm pore polystyrene membrane inserts (Corning, NY, USA), into which 1.4×10^10^ cells of *S. gordonii* or *V. parvula* individually, or their mixture (7×10^9^ cells each) in CDM, or an equal volume of fresh CDM (as a control) were added. Conditions were in triplicate and the setup was anaerobically incubated at 37°C. Anaerobic conditions were intended not only to reproduce ornithine cross-feeding (18), but to protect *F. nucleatum* from toxicity of H_2_O_2_ produced by *S. gordonii* in the presence of oxygen (49) which is unlikely to occur in the anaerobic microenvironment of the gingival margins and subgingival area, and to maximize the cooperative potential for metabolite exchange between these species. After 6 h, *F. nucleatum* cells were collected by pipetting from the lower chamber and washed with Milli-Q water by centrifugation. For metabolomics analysis, bacterial pellets were immediately fixed by adding methanol containing 5 μM internal standard. Spent medium from cultures and sterile CDM were centrifuged, filtered through 0.22-μm PES filtration devices (Merck Millipore, Darmstadt, Germany) and lyophilized. Capillary electrophoresis time-of-flight mass spectrometry (CE-TOFMS) was performed as described previously (18). As for metabolites whose levels were altered significantly in spent media of co-cultures, replicate experiments were performed with the additional control of mono-cultures of *S. gordonii* or *V. parvula* in CDM (1.4×10^10^ cells in the lower chamber of a Transwell plate), and metabolite concentrations in the culture supernatants were quantified using an Acquity ultra-performance liquid chromatography (UPLC) system with a PDA Detector (Waters, Milford, MA, USA), as described previously (18), with the exception of butyrate which was quantified using an high-performance liquid chromatography as described previously (50). Quantification of mRNA transcripts was performed by qRT-PCR as described previously (18). Primers are listed in Table S1. ATP was measured in a chemiluminescent assay as described previously (51). For additional details regarding CE-TOFMS, transcriptional analysis and ATP measurement, see Supplementary Materials and Methods.

### Polyamine production assay

Mono-cultures of *F. nucleatum* or *S. gordonii* (6.75×10^9^ cells), and mixed cultures of *F. nucleatum* with *S. gordonii* or *V. parvula* (6.75×10^9^ cells each) were anaerobically incubated at 37°C in pre-reduced CDM containing 10 mM arginine, or lysine. Culture supernatants were collected by centrifugation and filter sterilized with 0.22-μm PES filters (Merck Millipore). Metabolite concentrations in the culture supernatants were determined using an Acquity UPLC system with a PDA Detector (Waters), as described previously (18). pH values in the culture supernatants were determined with an F51 pH meter (Horiba, Kyoto, Japan).

### MALDI-MSI for putrescine visualization

MALDI-MSI was performed as described previously (52). Briefly, Indium-tin-oxide (ITO)-coated glass slides were immersed in *F. nucleatum* mono-cultures developed in PBS with or without 10 mM ornithine. After 24 h, biofilms formed on the glass slides were gently washed with PBS and subjected to fixation. After on-tissue derivatization using 2,4-diphenyl-pyranylium tetrafluoroborate (DPP-TFB, Merck, St. Louis, MO, USA), MALDI-MSI analyses were performed using iMScope TRIO (Shimadzu, Kyoto, Japan). See Supplementary Materials and Methods for more detail.

### Biofilm assay

To assess the effects of various polyamines on *P. gingivalis* biofilms, we initially preformed biofilms by incubating 4×10^7^ cells anaerobically at 37°C for 24 h in pre-reduced minimal medium (53) in a 25% saliva-coated well of an 8-well Chamber Slide System (Thermo Fisher Scientific, Waltham, MA, USA) with rotating. The resulting biofilms were incubated anaerobically in pre-reduced PBS containing each polyamine for 12 h. After staining with a Live/Dead BacLight kit (Molecular Probes, Eugene, OR, USA), gentle washing with PBS and extended anaerobic incubation in pre-reduced PBS containing each polyamine for 3 h, biofilm microstructures and newly released planktonic cells were observed with a Leica SP8 confocal laser scanning microscope (CLSM; Leica Microsystem, Wetzlar, Germany) and analyzed with Imaris 7.1.0 software (Bitplane, Belfast, UK), which is fully described in Supplementary Materials and Methods. For other bacteria, the same procedures were repeated, with the details for preforming the biofilms described in Supplementary Materials and Methods. To assess the effects of putrescine on biofilms, *S. gordonii* and *P. gingivalis* were stained with 15 mg/l hexidium iodide (HI; Thermo Fisher Scientific) and 4 mg/l 5-(and-6)-carboxyfluorescein and succinimidyl ester (FITC; Thermo Fisher Scientific), respectively. Preformed biofilms were treated with pre-reduced PBS containing putrescine for 12 h, followed by CLSM. To observe the responses of *P. gingivalis* to cell-free supernatants from 24-h cultures of *F. nucleatum* and *S. gordonii*, the culture supernatants were obtained by the same method as those of the polyamine production assays, with the pH adjusted to 7. *P. gingivalis* (2.8×10^8^ cells) was stained with FITC and incubated anaerobically for 24 h in pre-reduced PBS containing 50% cell-free supernatants, followed by CLSM. For analysis of mixed biofilm formation, *F. nucleatum* (3.2×10^8^ cells), stained with FITC, was anaerobically cultured in CDM for 24 h, washed gently by PBS, then co-cultured with 3.2×10^7^ cells of *P. gingivalis* labelled with DAPI, in the presence of ornithine for 24 h.

### Sample collection and detection of selected genes

We employed supragingival plaque samples, collected in our previous multi-omics study (45, 46), which was conducted from 2013 to 2014, with approval from the Osaka University Research Ethics Committee and in accordance with the principles of the Helsinki Declaration and STROBE guidelines for human observational studies. All participants provided written informed consent prior to enrolment. Detailed information about inclusion and exclusion criteria, oral examinations and sample collection are shown in Supplementary Materials and Methods. Characteristics of study participants are summarized in Table S2. Bacterial DNA was extracted using DNeasy PowerSoil Pro Kit (Qiagen, Hilden, Germany) protocol according to the manufacturer’s instructions. Primers and TaqMan probes (conjugated with FAM, ZEN and IBFQ) were designed based on the specific sequences for *arcD* of *S. gordonii*, FN0501 of *F. nucleatum*, 16S rRNA gene of *P. gingivalis* using nucleotide BLAST (NCBI), CLUSTALW (DDBJ) and PrimerQuest (Integrated DNA Technologies, Coralville, IA, USA). A universal probe/primer set previously described was designed with some modifications and used for standardization (54, 55). TaqMan real-time PCR was performed on a Roter-Gene Q System (Qiagen) using a Thunderbird SYBR qPCR Mix (Toyobo, Osaka, Japan). Primers and probes are listed in Table S3.

### Statistical analyses

Statistical analysis for intracellular metabolomics data was based on multivariate analysis by OPLS-DA using SIMCA-P software (version 14.0; Umetrics, Umeå, Sweden). Score plots and S-plots were constructed using Pareto scaling, and metabolites that contributed most to discrimination were chosen based on a p (corr) value >0.6. Statistical analysis for extracellular metabolomics data was based on comparison between groups with Mann-Whitney’s U test using SPSS (version 22; IBM, NY, USA). Extracellular levels of selected metabolites were compared between co-cultures and each mono-culture with one-way analysis of variance (ANOVA) followed by Tukey’s test using SPSS. The results from qRT-PCR and biofilm assays were analyzed by one-way ANOVA with post hoc paired comparison conducted with Dunnett’s test using SPSS. ROC curves and logistic regression were performed with R package (v4.0.3).

## Acknowledgments

We thank AMED-CREST for support through 18gm0710005h0206 (MK); MEXT/JSPS KAKENHI for support through 18H04068 (AA), 18H05387 (AA), 19H03862 (MK), and 18K17281 (AS); and NIH/NIDCR for support through DE012505, DE023193 and DE011111 (RJL). We are also thankful for excellent technical assistance from Miho Kakiuchi.

